# Clinical and genomic crosstalk between glucocorticoid receptor and estrogen receptor α in endometrial cancer

**DOI:** 10.1101/199448

**Authors:** Jeffery M. Vahrenkamp, Chieh-Hsiang Yang, Adriana C. Rodriguez, Aliyah Almomen, Kristofer C. Berrett, Alexis N. Trujillo, Katrin P. Guillen, Bryan E. Welm, Elke A. Jarboe, Margit M. Janat-Amsbury, Jason Gertz

## Abstract

Steroid hormone receptors are simultaneously active in many tissues and are capable of altering each other’s function. Estrogen receptor α (ER) and glucocorticoid receptor (GR) are expressed in the uterus and their ligands have opposing effects on uterine growth. In endometrial tumors with high ER expression, we surprisingly found that expression of GR is associated with poor prognosis. Dexamethasone reduced normal uterine growth *in vivo*; however, this growth inhibition was abolished in estrogen-induced endometrial hyperplasia. We observed low genomic binding site overlap when ER and GR are induced with their respective ligands; however, upon simultaneous induction they co-occupy more sites. GR binding is significantly altered by estradiol with GR recruited to ER bound loci that become more accessible upon estradiol induction. Gene expression responses to co-treatment were more similar to estradiol, but with novel regulated genes. Our results suggest phenotypic and molecular interplay between ER and GR in endometrial cancer.

## Introduction

Steroid hormone receptors have similar DNA binding preferences and their genomic binding is dependent on an overlapping set of additional transcription factors (Jozwik and Carroll, 2012; Robinson et al., 2011). Therefore, it isn’t surprising that the actions of one steroid hormone receptor can be altered by the induction of a different steroid hormone receptor. For example, progesterone receptor (PR) and estrogen receptor α (ER) are often co-expressed in breast cancer and the induction of PR redirects ER genomic binding and reduces estrogen-driven growth(Mohammed et al., 2015; Singhal et al., 2016). Androgens enable breast cancer growth and androgen receptor (AR) is important for ER genomic binding (D’Amato et al., 2016). Steroid hormones are also capable of compensating for one another. In prostate cancer, dexamethasone can confer resistance to anti-androgen therapy by activating glucocorticoid receptor (GR), which can substitute for AR in regulating transcription (Arora et al., 2013; Isikbay et al., 2014; Li et al., 2017). In the luminal androgen receptor subtype of breast cancer, AR compensates for ER’s absence by binding to very similar genomic loci that are also occupied by FOXA1(Robinson et al., 2011). It is clear that steroid hormone receptors do not work in isolation and that they can perform overlapping functions.

GR and ER have been shown to alter each other’s regulatory and phenotypic roles. GR expression is associated with good outcomes in ER-positive breast cancer and poor outcomes in ER-negative breast cancer (Pan et al., 2011b), which is consistent with dexamethasone blocking estrogen induced growth in breast cancer cells(Zhou et al., 1989). When both steroid hormone receptors are active in breast cancer cells, GR impacts ER’s genomic interactions (Miranda et al., 2013). GR activation alters chromatin accessibility enabling ER to bind to a new set of genomic regions, a mechanism known as assisted loading(Voss et al., 2011). In addition, ER and GR have been shown to both cooperate(Bolt et al., 2013) and compete(Karmakar et al., 2013; Meyer et al., 1989) for co-factors in breast cancer cells. GR has also been implicated in trans-repression of ER regulated gene expression in breast cancer (Yang et al., 2017). Estrogens and corticosteroids can elicit opposite phenotypic effects in other tissues as well(Haynes et al., 2003; Lam et al., 1996; Terakawa et al., 1985), including the uterus(Gunin et al., 2001; Markaverich et al., 1981; Rabin et al., 1990; Rhen et al., 2003).

Endometrial cancer is the most common gynecological cancer and incidence as well as mortality associated with endometrial cancer are on the rise, while survival is significantly worse now than in the 1970s. Of endometrial cancers, 80-90% are type I endometrioid tumors that express ER and are thought to be hormonally driven(Saso et al., 2011). Estrogen causes increased uterine growth and continued exposure can lead to endometrial hyperplasia(Yang et al., 2015). In contrast to ER’s pro-growth role in the uterus, GR reduces uterine growth and opposes phenotypic effects of estrogens in the uterus(Bever et al., 1956; Bitman and Cecil, 1967). Despite the opposing phenotypic roles, Dexamethasone (Dex) and 17β-estradiol (E2), a GR and an ER agonist respectively, produce similar gene expression changes in the normal uterus(Rhen et al., 2003). While estrogen and corticosteroid signaling have been examined in the normal uterus, crosstalk between ER and GR has not been explored in the context of endometrial cancer on a genome-wide scale.

Here, we show that GR expression is associated with poor prognosis and higher grade in endometrioid endometrial cancers, which is unexpected considering the growth inhibitory effects of corticosteroids in the normal uterus. Consistent with this observation, we find that once E2-induced hyperplasia is established in mice, growth is no longer opposed by Dex. Analysis of GR and ER genomic binding in endometrial cancer cells uncovered an interesting relationship in which E2, in combination with Dex, enables GR to bind to an expanded repertoire of genomic loci. Based on chromatin accessibility patterns, ER appears to assist in chromatin loading of GR. Gene expression analysis indicates that double induction with E2 and Dex produces a similar response as the addition of the separate inductions; however, several genes are impacted by the double treatment in a nonlinear manner. Together our results indicate that ER impacts GR’s gene regulatory and phenotypic actions in endometrial cancer cells and that GR activity may lead to more aggressive forms of the disease.

## Results

### Glucocorticoid Receptor expression is associated with poor prognosis in endometrial cancer

Because steroid hormone receptors can impact each other’s phenotypic and gene regulatory outputs, we decided to look for association between gene expression of steroid hormone receptors and outcomes in endometrioid endometrial cancer, a hormonally driven cancer type. We focused our analysis on endometrial cancer RNA-seq and clinical data from The Cancer Genome Atlas (TCGA) (Cancer Genome Atlas Research et al., 2013), exclusively analyzing tumors with endometrioid histology. While ER is expressed at the mRNA level in most tumors, higher expression is associated with longer disease-free survival (Figure 1A, p-value = 0.002791, Cox regression), which is consistent with previous reports for ER protein and mRNA levels(Wik et al., 2013). Higher ER expression was also associated with lower grade (Figure 1D, p-value = 1.75×10^−7^, Wilcoxon), as previously observed with ER protein levels(Backes et al., 2016), indicating that ER expression is higher in more differentiated tumors. PR is transcriptionally regulated by ER and therefore it isn’t surprising that higher expression of PR is also associated with better outcomes (Figure S1, p-value = 0.0006479, Cox regression) as previously reported(Tangen et al., 2014). Expression of AR, mineralocorticoid receptor, and estrogen receptor β were not significantly associated with recurrence (Figure S1).

**Figure 1.**
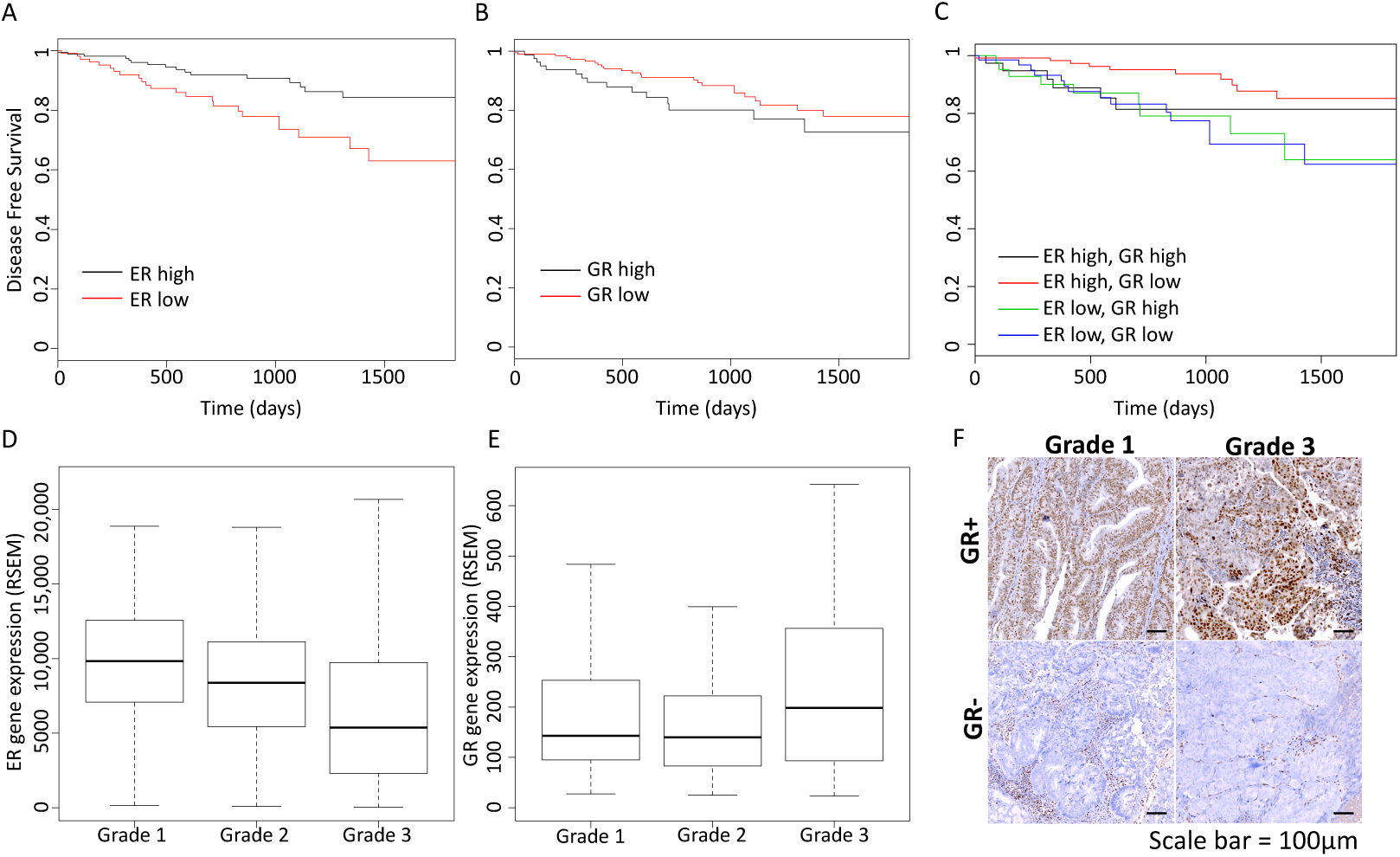
GR expression is associated with poor prognosis in endometrioid endometrial cancer. Kaplan-Meier curves show the association of disease-free survival with ER expression (A, high is top 60% tumors), GR expression (B, high is top 30% tumors), and the combination (C). Higher ER expression is associated with better prognosis, while higher GR expression is associated with worse outcomes, but only in the context of high ER expression. ER expression is negatively correlated with tumor grade (D) and GR expression is positively correlated with tumor grade (E). F) Examples of GR immunohistochemistry staining are shown for grade 1 and grade 3 tumors, scale bar represents 100 μm. See also Figures S1 and S2.

Higher GR expression was associated with worse outcomes for patients with endometrioid endometrial tumors (Figure 1B, p-value = 0.012, hazard ratio = 2.1, Cox regression). Higher expression was also correlated with higher histological grade (Figure 1E, p-value = 0.04, Wilcoxon). In order to assess protein expression and GR activity in endometrioid tumors, we performed IHC and found that 27% (3/11) of grade 1 endometrioid tumors exhibited nuclear GR staining, which is indicative of active GR, and 50% (5/10) of grade 3 endometrioid tumors were positive for nuclear GR staining (examples in Figure 1F, Figure S2). The IHC confirms that GR is expressed and active in endometrial cancer cells and is more likely to be present in higher grade endometrioid endometrial tumors.

To further explore the relationship between GR expression and poor prognosis, we analyzed this association in the higher ER expressing tumors and the lower ER expressing tumors separately. Figure 1C shows that association between GR expression and disease-free survival is only seen in the tumors with higher ER expression (p-value = 0.0096, hazard ratio = 3 [1.3 - 7.3 95% confidence interval], Cox regression) and not observed in the tumors with lower ER expression (p-value = 0.94, hazard ratio = 0.97 [0.42 - 2.23 95% confidence interval], Cox regression). These results suggest that GR expression increases the aggressiveness of tumors with higher expression of ER, but has little effect on tumors with low ER expression indicating that ER and GR could be impacting the actions of one another.

### Estradiol and dexamethasone in combination induce growth of endometrial cancer cells in 3D culture

Positive correlation between GR expression and the likelihood of recurrence is surprising, and appears inconsistent, in light of multiple studies connecting GR activation with reduced uterine growth (Gunin et al., 2001; Markaverich et al., 1981; Rabin et al., 1990; Rhen et al., 2003). We hypothesized that the growth inhibitory effects of corticosteroids may be absent once hyperplasia is established. We first evaluated how ER and GR affect endometrial cancer cell growth *in vitro*. The endometrial cancer cell line Ishikawa was grown in 3D matrigel culture in the presence of hormone depleted media and then treated with E2, Dex or the combination for 14 days. We then measured ATP levels and observed an expected loss of cell growth in hormone depleted media (Figure 2A). The addition of E2 and Dex individually had subtle and insignificant effects on proliferation. However, when organoids were treated with both E2 and Dex, there was a significant increase in growth that came close to full media levels. This evidence indicates that ER and GR work together in promoting the growth of endometrial cancer cells.

**Figure 2.**
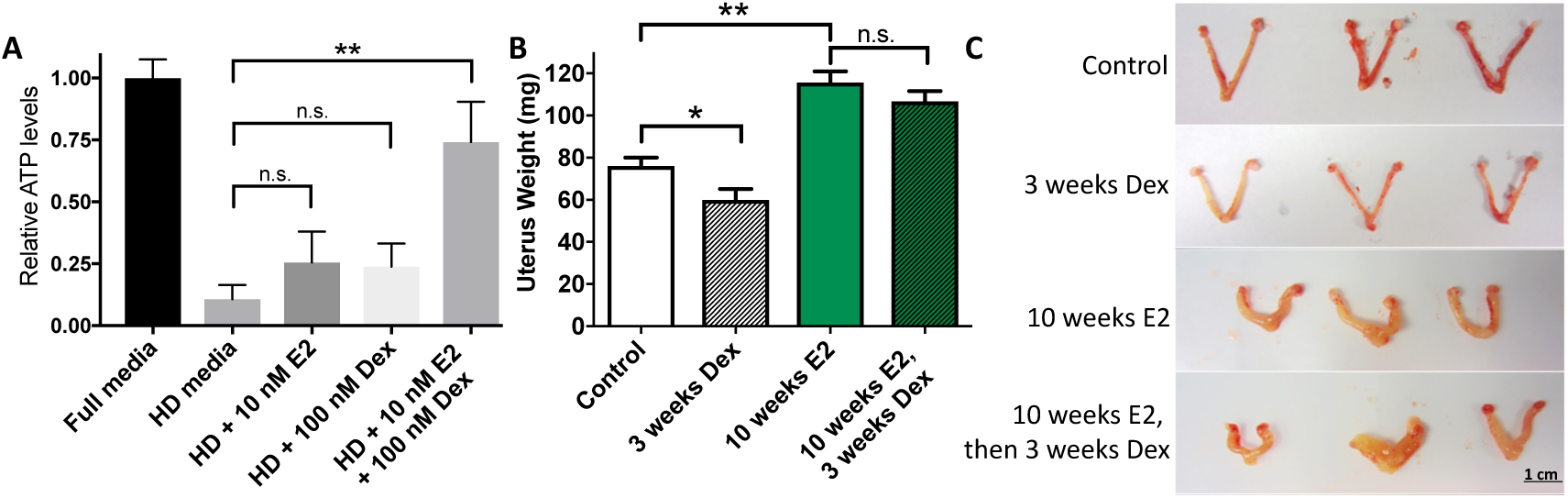
Growth effects of dexamethasone are impacted by estradiol. A) Hormone depleted (HD) media reduces growth of Ishikawa cells, as measured by relative ATP levels. The addition of E2 or Dex to HD media modestly increases growth and the combination of E2 and Dex restores growth to similar levels as full media. B) Uterine weight is reduced by Dex in mice. Continuous E2 exposure for 10 weeks significantly increases uterine weight and subsequent addition of Dex does not significantly reduce this effect. C) Examples of uteri from the categories in panel B, scale bar represents 1 cm. Data are represented as mean ± SEM in panels A and B. * = p-value < 0.05, ** = p-value < 0.01, and n.s. = not significant (p-value > 0.05). See also Figure S3.

### Dexamethasone inhibition of uterine growth is not observed once estrogen-driven hyperplasia is established

We next sought to determine the impact of active ER and GR on endometrial cells *in vivo*. We used an estrogen-induced endometrial hyperplasia model that utilizes a slow release E2 pellet(Yang et al., 2015). We found that 3 weeks of Dex treatment, in the absence of excess E2, reduced uterine weight by 21% (p-value = 0.042, t-test), while 10 weeks of E2 exposure increased uterine weight by 52% (p-value = 0.002, t-test) (Figure 2B,C). When we administered Dex after allowing hyperplasia to establish during 10 weeks of excess E2, we found that Dex was unable to significantly inhibit uterine growth (p-value = 0.24, t-test; 7.8% decrease) (Figure 2B,C). Histo-pathological review of hematoxylin & eosin (H&E) stained slides from harvested and processed uteri revealed that endometrial hyperplasia persisted in 5 out of 6 mice treated with Dex after E2, as compared to 7 out of 7 mice treated with only E2. We also observed the largest effects on endometrium growth, which is significantly thicker after E2 treatment (Figure S3; p-value = 6.7×10^−6^, Wilcoxon), while myometrium growth is slightly reduced by E2 (p-value = 0.0179, Wilcoxon). Overall, these results suggest that GR’s inhibitory effect on uterine growth is significantly reduced once hyperplasia is established.

### Glucocorticoid Receptor binds loci occupied by ER upon estradiol and dexamethasone treatment

The survival analysis and phenotypic observations suggest that ER and GR may be impacting the role of one another. In order to determine how this crosstalk is occurring at a molecular level, we performed ChIP-seq with antibodies targeting ER and GR in Ishikawa, a human endometrial adenocarcinoma cell line. We found that ER and GR, when induced by E2 and Dex respectively, have mostly unique binding profiles with only 19.7% of GR bound loci overlapping ER bound loci. However, when cells are treated with E2 and Dex simultaneously, GR and ER share almost half of their binding sites (46.5%). This co-occurrence in ER and GR binding upon the double induction represents a significant increase compared to ER and GR after single inductions (p-value < 2.2×10^−16^, odds ratio = 3.96, Fisher’s Exact Test, all overlaps shown in Figure S4A). These findings indicate that ER and GR are impacting one another on the level of genomic binding site selection.

Further analysis of the ER and GR bound sites following double induction revealed that GR is moving to sites that are bound by ER. The vast majority (91.8%) of ER bound sites in the double induction were bound by ER in cells treated with E2 alone (Figure 3A) and the sites unique to the double induction appear to be accompanied by small changes in ChIP-seq signal (Figure 3B). In contrast, 63.1% of GR bound sites in the double induction were bound by GR in cells treated only with Dex and the sites unique to the double induction exhibit large changes in ChIP-seq signal (Figure 3C). Of the 598 sites that were bound by GR only in the double induction, 67.5% overlapped with ER suggesting that most of the new GR binding sites in the double induction are loci that are bound by ER after both E2 and E2 + Dex treatments. The presence of estrogen response elements (EREs) and glucocorticoid receptor binding elements (GRBEs) are consistent with this idea; we found that loci bound by GR only after the double induction are enriched for both full and half site EREs compared to GRBEs (Table 1). The increase in GR genomic binding is not due to increased expression as GR mRNA and protein levels are unaffected by the treatments (Figure S4B,C). Overall, these results indicate that ER is enabling GR to bind new genomic loci when both steroid hormone receptors are active.

**Figure 3.**
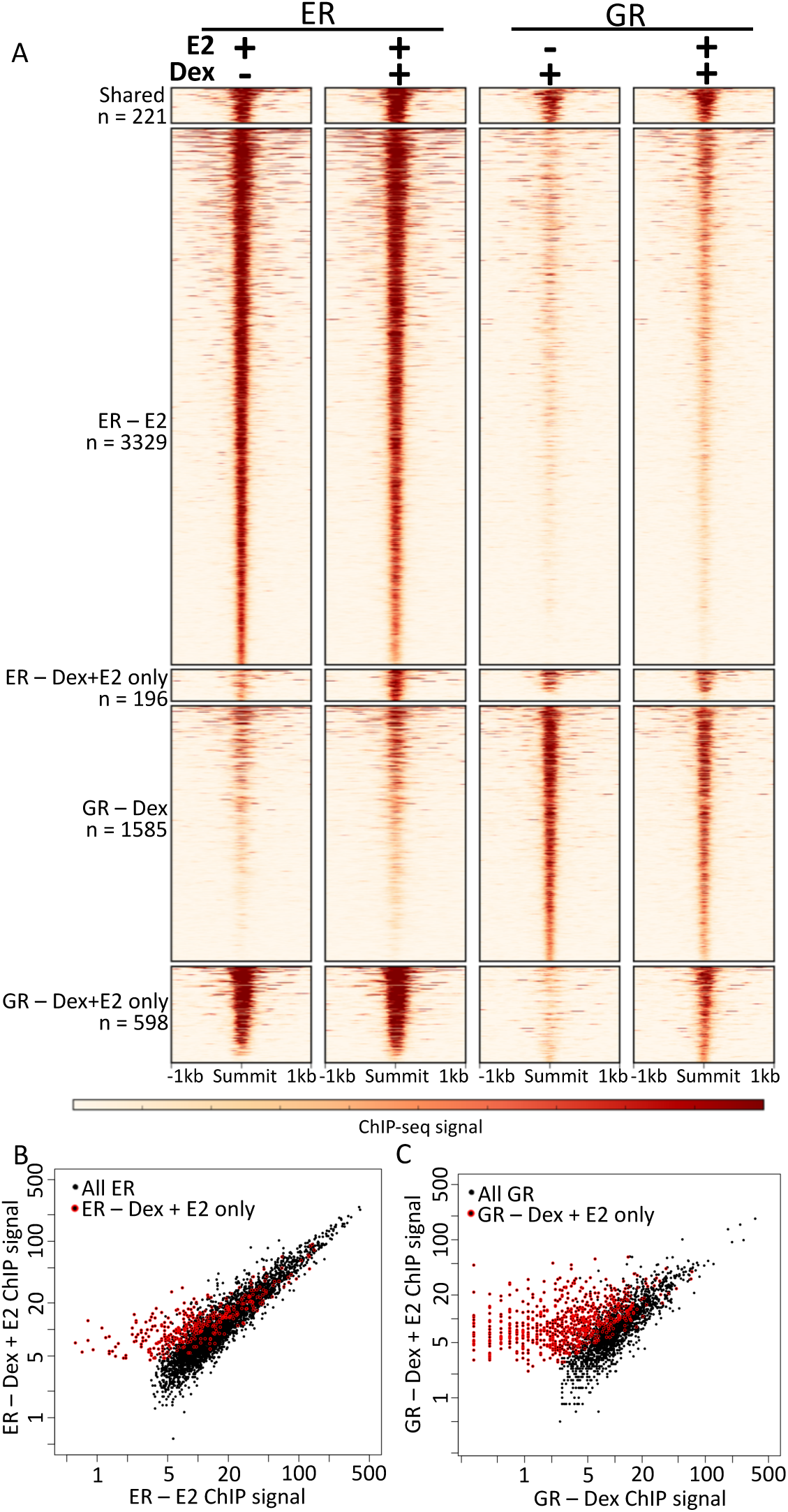
Estradiol alters GR genomic binding. A) Heatmaps of ChIP-seq signal of ER and GR after single induction and double inductions show that there are a unique set of sites where GR binds only after double induction. ChIP-seq signal is plotted for each ER and GR (C) binding site comparing signal induction (x-axis) and double induction (y-axis). Sites unique to the double induction are shown in red with GR double induction sites showing markedly different signal. Signal is displayed as reads per million. See also Figure S4.

**Table 1.** Motif occurrences at ER and GR bound loci.

| Motif | ER – E2 | ER – Dex + E2 only | GR – Dex | GR – Dex + E2 only |
| --- | --- | --- | --- | --- |
| ERE full site | 27% | 8% | 3% | 21% |
| ERE half site | 91% | 76% | 55% | 82% |
| GRBE full site | 6% | 14% | 45% | 11% |
| GRBE half site | 53% | 64% | 81% | 60% |

### Histone acetylation is increased by estradiol and dexamethasone co-treatment at sites bound by ER and GR

To determine if binding of ER and GR to genomic loci upon double induction impacted the regulatory activity of these regions, we analyzed acetylation of histone H3 lysine 27 (H3K27ac), a mark associated with active regulatory regions. We performed ChIP-seq, with an antibody that recognizes H3K27ac, on cells that were treated with Dex, E2, the combination, or DMSO (vehicle control) for 8 hours. We found that at GR sites bound in the Dex only treatment there were not significant changes in H3K27ac after any treatment in comparison to vehicle (Figure 4A; p-values > 0.01, Wilcoxon). Sites bound by ER after E2 only treatment exhibited higher levels of H3K27ac after E2 treatment and the combination treatment compared to vehicle (Figure 4B; p-values < 2.2×10^−16^, Wilcoxon). Genomic loci that are bound by both GR and ER following the combination treatment had significantly higher levels of H3K27ac after the combination treatment compared to vehicle, E2 alone, and Dex alone (Figure 4C; p-value = 2.3×10^−5^, Wilcoxon). The pattern of H3K27ac enrichment at loci bound by both ER and GR indicate that binding of ER and GR to the same locus leads to molecular changes that are not observed when only one factor is active.

**Figure 4.**
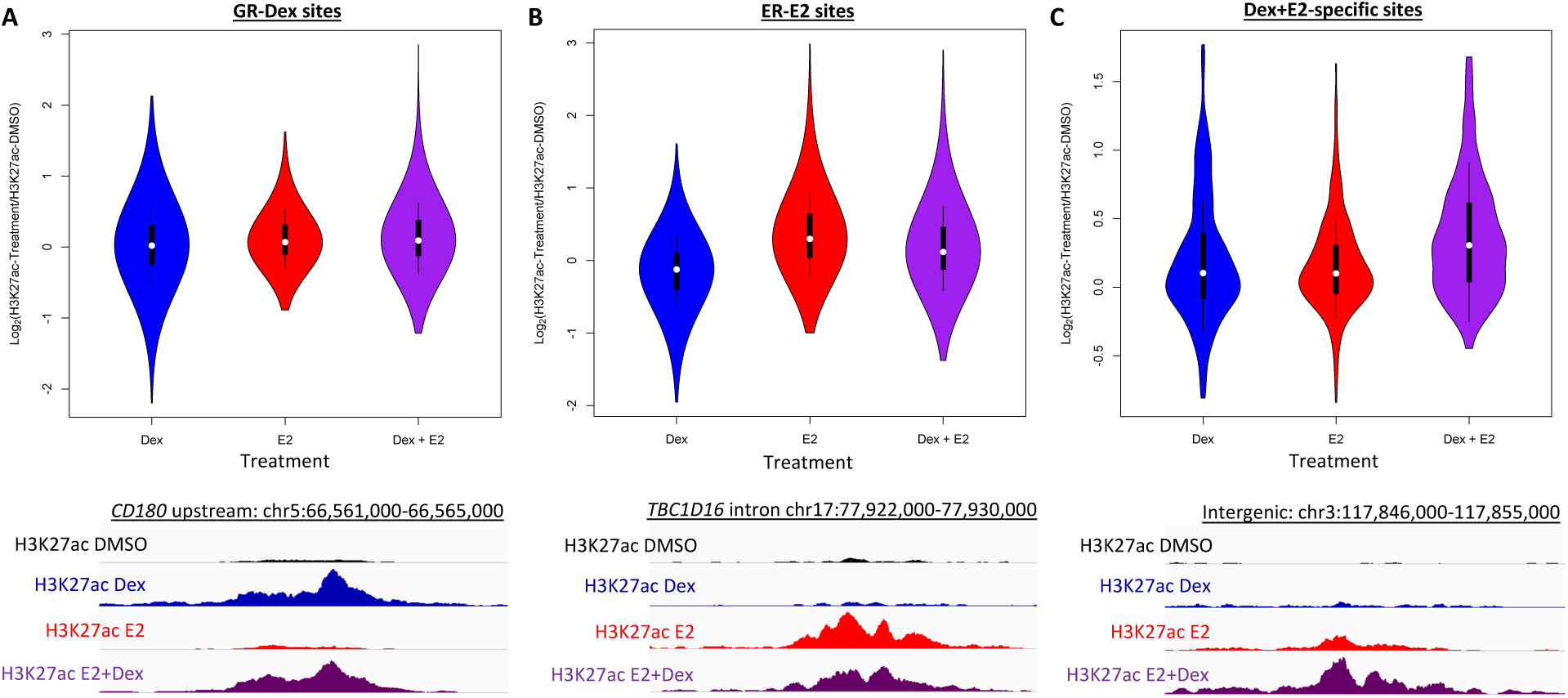
Histone acetylation is increased at ER and GR bound sites upon double induction. For GR sites bound after Dex only treatment (A), ER sites bound after E2 only treatment (B), and sites bound by both ER and GR after double induction (C), the distribution of relative levels of H3K27ac for Dex (blue), E2 (red), and the combination (purple) compared to the vehicle control are shown. H3K27ac increases at loci bound by both ER and GR specifically after the combination treatment. Example loci for each type of site are shown in the bottom panels with each track for a site shown on the same scale.

### Chromatin accessibility patterns are consistent with ER assisting GR in chromatin loading

One potential explanation for ER enabling GR genomic binding is assisted loading, where one transcription factor makes it easier for another factor to bind by increasing accessibility to the genomic site(Voss et al., 2011). This phenomenon has been observed in breast cancer cells, although in the opposite direction with GR enabling ER binding (Miranda et al., 2013). To test this model, we performed ATAC-seq on Ishikawa cells treated with Dex, E2 or vehicle controls for 1 hour. We focused our analysis first on loci that were bound by GR either after induction with Dex or only after induction with both E2 and Dex. The loci bound only after the double induction exhibited significantly higher E2-induced chromatin accessibility as measured by ATAC-seq (Figure 5A; p-value< 2.2×10^−16^, Wilcoxon). This result is consistent with ER creating a more permissive binding environment for GR, enabling GR to bind only after the double induction. Figure 5C and 5D show example loci bound by GR only after double induction that also become more accessible upon E2 induction. In total, 65% of loci bound by GR only after the double induction exhibit increased accessibility after E2 treatment, compared to 43% of all GR bound loci. We performed a similar analysis for loci bound by ER only after double induction and found that chromatin accessibility significantly increases upon Dex induction (Figure 5B; p-value = 0.0001223, Wilcoxon), but the effect is more subtle with 55% of these loci showing increased accessibility after Dex treatment. The ATAC-seq findings are consistent with a model where ER increases chromatin accessibility, allowing GR to bind new loci.

**Figure 5.**
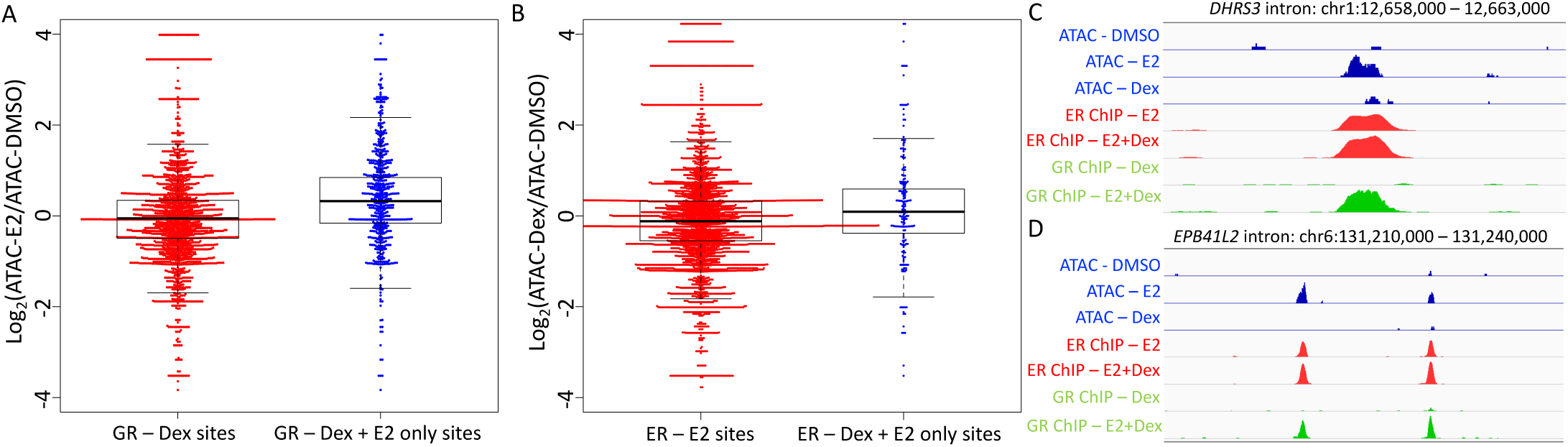
Chromatin accessibility is increased at double induction-specific GR bound sites. The log ratio of ATAC-seq signal after E2 treatment vs. vehicle control (A) and Dex treatment vs. vehicle controls (B) is shown for all (red) and double induction-specific (blue) GR (A) and ER (B) sites. ATAC-seq signal is increased upon E2 induction at GR sites only observed after E2 + Dex treatment. Signal tracks of ATAC-seq signal (blue), ER ChIP-seq signal (red) and GR ChIP-seq signal (green) are shown for loci nearby *DHRS3* and *EPB4IL2* (D). Signal tracks of the same type are shown on the same scale. See also Figure S6.

### The combination of estradiol and dexamethasone cooperate to regulate gene expression

We sought to determine how crosstalk between ER and GR at the genomic binding level impacts gene expression. In order to analyze gene expression responses to E2, Dex and the combination, we performed RNA-seq and analyzed differential gene expression. We identified 286 genes that were regulated by E2 alone (188 up-regulated and 98 down-regulated) and 128 genes significantly regulated by Dex alone (120 up-regulated and 8 down-regulated). There was little overlap in the genes that were regulated by both E2 and Dex individual treatments with 24 overlapping genes that go in the same direction (19% of all Dex regulated genes) and 6 overlapping genes that are regulated in opposite directions (5% of all Dex regulated genes) (Figure S5A). *PR*, which is growth inhibitory in the uterus, is an example of an independent gene that is unaffected by Dex and induced by E2 and the combination treatment to a similar level. We examined other growth inhibitory genes, including *EIG121, RALDH2, SFRP1*, and *SFRP4* (Deng et al., 2010; Westin et al., 2009), and found that each gene exhibited very low expression in Ishikawa cells.

Upon treatment with both E2 and Dex, 371 genes were significantly affected (240 up-regulated and 131 down-regulated) and the combination treatment samples appeared more similar to the E2 treated samples. Most E2 induced gene expression changes were observed with the combination (69%, Figure S5B), while less than half of the Dex induced expression changes were also seen with the combination treatment (43%, Figure S5C). This relationship is also clear when analyzing principal components (Figure 6A). The overall pattern of gene expression upon E2 and Dex treatment suggests that E2 dominates the combination treatment, explaining the pro-growth phenotype, but Dex contributes by expanding the genes affected. The genes unique to the double induction are enriched in down-regulation of cell-cell adherens junctions (adjusted p-value = 0.001, DAVID(Jiao et al., 2012)), including alpha catenin(Knudsen, 1995), vinculin(Carisey and Ballestrem, 2011) and cingulin(Citi et al., 1988) (Figure S5D-F). The down regulation of cell adhesion proteins could partially explain the aggressiveness of ER and GR expressing tumors (Figure 1C). The genes uniquely regulated by the combination treatment are also found nearby genomic loci bound by ER and GR after the double induction. GR double induction sites are significantly closer to the Dex + E2-specific genes than single induction GR sites (Figure S5G, p-value = 4.811×10^−4^, Wilcoxon). ER double induction sites are also closer to Dex + E2-specific genes, but the effect is more subtle and the p-value is marginal (Figure S5H, p-value = 0.0421, Wilcoxon). These findings are consistent with a shift in GR sites upon double induction leading to novel regulation in cells co-treated with E2 and Dex.

**Figure 6.**
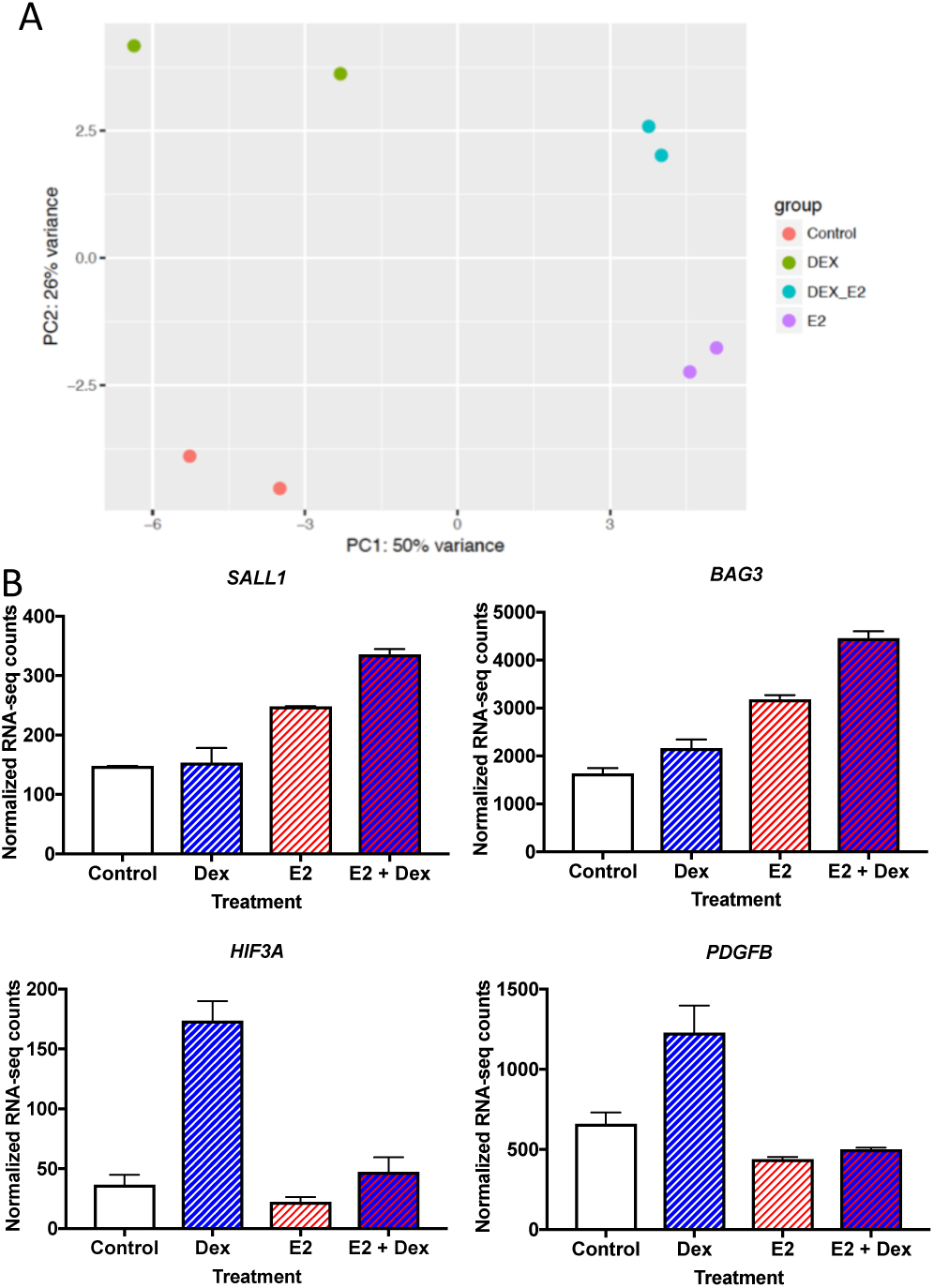
Gene expression consequences of double induction. A) Principal component analysis of RNA-seq from E2, Dex and the combination treatments, shows that E2+Dex (cyan) treated samples are more similar to E2 (purple) treated samples than Dex treated (green) or vehicle treated (red) samples. B) Examples of genes that exhibit unexpected gene expression levels upon double treatment are *SALL1, BAG3, HIF3A*, and *PDGFB*. Data are represented as mean ± SEM. See also Figures S5 and S7.

For the 525 genes that exhibited differential gene expression in any treatment compared to controls, we performed regression analysis to look for genes that have a significant E2:Dex interaction term. We identified 112 genes with significant interactions terms; 21 genes had higher expression in the double induction than expected based on the single inductions and 91 genes had lower than expected expression in the double induction. Of the genes with lower than expected double induction expression, 61 (67%) were affected more by Dex alone than E2 alone, indicating a loss of Dex responsiveness that is consistent with altered GR binding. Figure 6B shows the expression of *HIF3A* and *PDGFB* as examples. Both *HIF3A* and *PDGFB* have regulatory loci nearby that exhibit less ATAC-seq signal upon E2 treatment as well as less GR binding after the double induction (Figure S6A, B). Of the genes with higher than expected double induction expression, 62% were affected more by E2 alone than Dex alone, suggesting that GR may be assisting ER in regulating some genes. The examples of *SALL1* and *BAG3* are shown in Figure 6B. The intron of *BAG3* exhibits increased ER binding with the double induction and a *SALL1* downstream site is only bound by GR upon treatment with both E2 and Dex (Figure S6C, D). Our results indicate that ER and GR can influence one another’s ability to regulate gene expression through both increasing and decreasing genomic binding.

### Many genes regulated by dexamethasone and estradiol are differentially expressed in endometrial tumors

In order to determine if the gene expression changes we observed in Ishikawa cells are also observed in patient tumors, we analyzed gene expression from the endometrioid histology endometrial tumors in the TCGA cohort. We first categorized tumors as having high or low mRNA expression of ER and GR. For each gene, we then compared gene expression in the GR high-ER high tumors to all other tumors. Of the genes that are differentially expressed when treated with the combination of Dex and E2 in Ishikawa cells, 37% were also differentially expressed in the GR high-ER high tumors, which represents a highly significant overlap (p-value = 1.285×10^−6^, Fisher’s exact test). Several examples of these genes are shown in Figure S7, which includes alpha catenin which is discussed above. We also found that nine of these genes exhibit expression that is associated with disease-free survival in endometrial cancer (Figure S7, *LMCD1* not shown due to space). These findings show that many of the genes uniquely regulated by Dex and E2 in Ishikawa cells are also differentially expressed in GR high-ER high endometrial tumors, indicating that ER and GR are likely to exhibit molecular crosstalk in patient tumors.

## Discussion

ER and GR play opposite phenotypic roles in the normal endometrium with ER promoting growth and GR inhibiting growth. Here we show that in endometrial cancer, GR expression is associated with worse outcomes, as well as higher grade tumors, and this association is only observed in the context of high ER expression. These findings seem to contradict the anti-growth effects of corticosteroids; however, our results show that growth inhibition is no longer observed after estrogen-induced hyperplasia has formed *in vivo* and that Dex, in combination with E2, promotes endometrial cancer cell growth in culture.

One possible explanation for the difference in GR induced growth effects is that ER and GR are expressed in different compartments of the endometrium with GR expressed in stromal cells and ER is expressed in endometrial glands(Bamberger et al., 2001). GR signaling in stromal cells, through either autocrine or paracrine changes, causes growth inhibition of the normal uterus. However, GR activity in hyperplastic or cancerous endometrial cells, when co-expressed in the same cells as ER, no longer inhibits growth and may lead to more aggressive tumors. In fact, gene expression profiling of cells induced with both Dex and E2 uncovered a down regulation of cell adherence genes, which has the potential to cause a more metastatic phenotype. These findings are consistent with work in breast cancer showing that GR expressing tumors differentially regulate cell adhesion genes(Pan et al., 2011a). Our findings indicate that the administration of dexamethasone during treatment of endometrioid histology endometrial cancer should be re-evaluated.

Molecular characterization of ER and GR crosstalk in endometrial cancer cells revealed that ER is the dominant steroid hormone receptor in this setting. ER genomic binding is mostly unaffected by the activation of GR. On the other hand, more than one third of GR binding is altered by the activation of ER. It also appears that recruitment of GR to ER bound sites has a functional consequence as it likely leads to an increase in regulatory activity as measured by H3K27ac and proximity to regulated genes. ER dominance is the opposite pattern than is observed in breast cancer cells, where GR is dominant in dictating ER binding (Miranda et al., 2013). Interestingly, the difference in dominance between GR and ER is associated with differences in prognosis. GR is dominant over ER in breast cancer cells resulting in a better prognosis when both are expressed (Pan et al., 2011b), while ER is dominant over GR in endometrial cancer cells resulting in a worse prognosis when both are expressed (Figure 1C). This leads to a model in which GR is generally growth inhibitory, ER is generally growth promoting, and the dominant factor dictates the aggressiveness of the tumor. It is unclear why ER is dominant in one setting and GR is dominant in another, but differences in co-factor abundance is an interesting possibility.

To explore how ER was altering GR genomic binding, we used ATAC-seq to identify changes in chromatin accessibility. We found that the majority of GR binding sites that are gained upon treatment with both Dex and E2 exhibit increased chromatin accessibility after E2 induction. These results are consistent with ER assisting in loading GR onto genomic loci, which is similar to breast cancer cells where GR assists in loading ER onto genomic loci (Miranda et al., 2013). We attempted co-immunoprecipitation of ER and GR and were unable to pull down one factor with an antibody that recognizes the other factor (data not shown), suggesting that ER and GR are not forming heterodimers while further supporting the model that crosstalk between ER and GR is occurring through alteration of each other’s chromatin interactions.

The genomic binding crosstalk between ER and GR has gene expression consequences. When endometrial cancer cells were induced with either Dex or E2 in isolation, there was very little overlap in the genes affected. When cells were treated with both Dex and E2, the transcriptional response was more similar to an E2 response, but with additional genes changing expression, including a down regulation of cell-cell adherens junction genes. A significant fraction of these additional genes are also differentially expressed in ER high-GR high endometrial tumors, indicating that many genes are regulated by the combination of ER and GR generally in endometrial cancer. Unlike a recent report in breast cancer(Yang et al., 2017), GR does not appear to repress an E2 transcriptional response in endometrial cancer cells. Overall, about one-fifth of genes affected by any treatment exhibited an unexpected gene expression level after double induction as determined by significant interaction terms in a linear model. Taken together, our findings are consistent with ER altering GR’s genomic actions, where GR switches from regulating a distinct set of genes to promoting and enhancing an E2-driven transcriptional program in endometrial cells.

## Experimental procedures

### TCGA data analysis

RNA-seq and clinical data was downloaded from the TCGA data portal in December 2015. Gene expression measurements were taken from the level 3 RNAseqV2 normalized RSEM data. Only samples with Endometrioid histology were analyzed for survival analysis. Cox regression to evaluate the association between gene expression and progression-free survival was performed in R using binary classification of high and low expression for each steroid hormone receptor. For each steroid hormone receptor, we tried each decade percentile increment between 20th percentile and 80th percentile in order to identify the cutoff between high and low expression that gave the most significant association with disease-free survival. The following cutoffs were used for each gene: GR - top 30th percentile was labelled high, ER - top 60th percentile, PR - top 80th percentile, AR - top 20th percentile, mineralocorticoid receptor - top 60th percentile, and estrogen receptor β - top 60th percentile. In order to identify genes that are differentially expressed in GR high-ER high tumors, we used a median cutoff for each gene to classify tumors as high or low. We then performed a Wilcoxon rank sum test to compare the expression in GR high-ER high tumors to all other endometrioid histology tumors.

### Immunohistochemistry

Immunohistochemical staining of formalin-fixed paraffin slides was performed by ARUP laboratories. In brief, 4-5 um thick tissue sections were prepared. The slides were placed on the automated immunostainer and de-paraffinized with the EZ Prep solution. The slides were then treated with CC1 (Cell Conditioning 1, pH 8.5) for 68 minutes at 95°C. The primary antibody against GR (Cell Signaling, clone D6H2L, #12041) was applied for 1 hour at a dilution of 1:100 at 35°C. Following removal of the primary antibody, the secondary antibody (Goat anti-rabbit IgG Sigma-aldrich, # B8895) was applied for 1 hour at a dilution of 1:100 at 37°C. These slices were then exposed to the IView DAB Map detection kit and counterstained with hematoxylin for 8 minutes. Slides were removed from the autostainer and placed in a dH2O/DAWN mixture, and then dehydrated in graded alcohol. After dipping each slide 10 times in 4 changes of xylene, cover slips were put on and slides were read by board certified pathologists. For patient samples, each patient was consented under a protocol approved by the Institutional Review Board of the University of Utah. All patients were female and age information was not analyzed for these patients.

### Animal studies

A total of thirty-five 8-week-old female Balb/c mice were used in this study. To induce estrogen-driven endometrial hyperplasia, E2 pellets were prepared and surgically implanted into mice (n=21) as previously described (Yang et al., 2015). Following 10 weeks of E2 exposure, mice were divided into three treatment arms to receive Dex, PBS, and 5% ethanol (n=7 per arm). Fourteen mice did not receive E2 pellet implants and were divided into two groups for administration of Dex and PBS (n=7 per group). Dex was dissolved in 5% ethanol and administered through intraperitoneal injection at a dose of 1mg/kg and a frequency of 5 days on 2 days off for a total of 21 days. Animals were sacrificed upon reaching the primary study endpoint on day 22 and necropsy including tissue harvesting ensued. Uteri were grossly examined and uterine weights were measured. Tissues were processed into FFPE blocks and stained with hematoxylin and eosin for histo-pathological analysis. Animals were housed in the Animal Facility of the Comparative Medicine Center at the University of Utah under standard conditions. All procedures conducted were approved by the Institutional Animal Care and Use Committee of the University of Utah.

### Cell culture

Ishikawa cells (Sigma-Aldrich) were grown in RPMI-1640 with 10% fetal bovine serum and 50 units/mL penicillin and 50 µg/mL streptomycin. At least four days before induction, cells were moved to phenol-red free RPMI-1640 with 10% charcoal-dextran stripped fetal bovine serum to remove hormones. Cells were treated with DMSO, 10 nM E2, 100 nM Dex or the combination for either 1 hour for ChIP-seq and ATAC-seq or 8 hours for RNA-seq.

### 3D Culture

Ishikawa cells were grown and transferred to hormone-depleted RPMI-1640 as described above. Cells were suspended in Matrigel (Corning, growth factor reduced, phenol-red free) and 500 cells were plated with 9 µL Matrigel and 500 µL media per well in a 48-well glass bottomed plate (MalTek, 48mm glass bottom). Cells were treated with DMSO, 10nM E2, 100nM Dex or combination of both 10nM E2 and 100nM Dex. Relative ATP levels were measured using CellTiter Glo 3D (Promega).

### ChIP-seq

After inductions, cells were crosslinked with 1% formaldehyde for 10 minutes at room temperature and then treated with 125 mM Glycine for 5 minutes to stop crosslinking. Crosslinked cells were then washed with cold PBS and scraped to harvest. Chromatin immunoprecipitation was performed as previously described (Reddy et al., 2009). Sonication was performed on an Active Motif EpiShear probe-in sonicator with 8 cycles of 30 seconds, with 30 seconds of rest, at 40% amplitude. The antibodies used were ER (Santa Cruz HC-20), GR (Santa Cruz E-20), and H3K27ac (Active Motif 39133). ChIP-seq reads for each treatment were compared to ChIP-seq libraries from control (DMSO) treated cells for the same factor (e.g. GR in Dex was compared to GR in DMSO). Reads were aligned to the hg19 build of the human genome using Bowtie with the following parameters: -m 1 -t --best -q -S -l 32 -e 80 -n 2. Peaks were called using MACS2(Zhang et al., 2008a) with a p-value cutoff of 1e-10 and the mfold parameter constrained between 15 and 100. Counts between sets were calculated using bedtools coverage(Quinlan and Hall, 2010) to find read depth at all sites covered by peaks called by MACS2. These data were combined using gawk and examined and graphed using R. Motif finding was performed on 100bp surrounding the top 500 peaks based on their integer score (column 5 of narrowPeak file). Motifs were discovered using the meme suite(Bailey et al., 2009) searching for motifs between 6 and 50 bases in length with zero or one occurrence per sequence. We used Patser(Hertz and Stormo, 1999) to count the number of sites with significant full and half site EREs and GRBEs within 50bp of each peak’s summit.

### RNA-seq

After inductions, cell lysates were harvested with buffer RLT Plus (Qiagen) supplemented with 1% beta-mercaptoethanol and passed through a 21-gauge needle to shear genomic DNA. RNA was purified using RNA clean and concentrator with the optional DNase treatment (Zymo Research). Poly(A)-selected RNA-seq libraries were constructed with KAPA Stranded mRNA-seq kit (Kapa Biosystems) using 1 ug total RNA. Sequencing reads were aligned to the hg19 build of the human genome using HISAT2 (Kim et al., 2015). Sam files were converted to bam format and sorted using samtools (Li et al., 2009). Reads mapping to genes were counted using featuresCounts from the SubRead package(Liao et al., 2014). Reads were normalized and differential analysis was done in a pairwise manner using DESeq2 (Love et al., 2014). Genes were considered significant if they had an adjusted p-value of <= 0.1. In order to find significant E2:Dex interaction terms, we constructed linear models in R using the ‘lm’ function and looked for E2:Dex coefficients that are significantly different than 0 with a p-value<0.05.

### ATAC-seq

ATAC-seq was performed on 250,000 cells for each library as described by Buenorostro et al (Buenrostro et al., 2013). Tn5 transposase, with Illumina adapters, was constructed as outlined by Picelli et al (Picelli et al., 2014). Sequencing reads were aligned to hg19 using bowtie(Langmead et al., 2009) with the following parameters: : -m 1 -t --best -q -S -l 32 -e 80 -n 2. Sam files were converted to bam files sorted using samtools (Li et al., 2009). Macs2 (Zhang et al., 2008b) was used to call peaks without a control input. We used a cutoff p-value of 1e-10 when calling peaks. FeatureCounts (Liao et al., 2014) was used to quantify reads that aligned in regions +/- 250 bp from an ER or GR binding site summits from the ChIP-seq experiments. These reads were then normalized and differential expression was determined by comparing samples in a pairwise manner using the DESeq2 package for R (Love et al., 2014).

### Statistical Analysis

All statistical analyses were performed in R version 3.4.0, with the exception of the p-value calculated by DAVID, and the statistical test used for each analysis is listed next to the reported p-value. The R functions used were wilcox.test, t.test, fisher.test, and coxph (from the package survival).

## Accession Numbers

The ChIP-seq, RNA-seq, and ATAC-seq data that was collected as part of this study is available at the Gene Expression Omnibus (GEO) under accession number GSE109893. Alignments, peak calls, and gene read counts, are included in the GEO submission.

## Acknowledgements

This work was supported by NIH/NHGRI R00 HG006922 and NIH/NHGRI R01 HG008974 to J.G., the Huntsman Cancer Institute, and the Women’s Cancers Disease-Oriented Team at the Huntsman Cancer Institute. Research reported in this publication utilized the High-Throughput Genomics Shared Resource at the University of Utah and was supported by NIH/NCI award P30 CA042014. A.C.R. was supported by diversity supplement R00 HG006922S1. We thank Ed Grow for providing reagents and we thank K-T Varley as well as Gertz and Varley lab members for their helpful comments on the study and the manuscript.

## Author contributions

Conceptualization, J.M.V., M.M.J., and J.G.; Investigation, J.M.V., C.Y., A.C.R., A.A., K.B., A.N.T., K.P.G., and E.A.J; Supervision, B.E.W., E.A.J., M.M.J., and J.G.; Writing – Original Draft, J.M.V., A.C.R., and J.G.; Writing – Review & Editing, all authors; Funding Acquisition, J.G.

## References

Arora, V.K., Schenkein, E., Murali, R., Subudhi, S.K., Wongvipat, J., Balbas, M.D., Shah, N., Cai, L., Efstathiou, E., Logothetis, C., et al. (2013). Glucocorticoid receptor confers resistance to antiandrogens by bypassing androgen receptor blockade. Cell 155, 1309–1322.

Backes, F.J., Walker, C.J., Goodfellow, P.J., Hade, E.M., Agarwal, G., Mutch, D., Cohn, D.E., and Suarez, A.A. (2016). Estrogen receptor-alpha as a predictive biomarker in endometrioid endometrial cancer. Gynecol Oncol 141, 312–317.

Bailey, T.L., Boden, M., Buske, F.A., Frith, M., Grant, C.E., Clementi, L., Ren, J., Li, W.W., and Noble, W.S. (2009). MEME SUITE: tools for motif discovery and searching. Nucleic Acids Res 37, W202–208.

Bamberger, A.M., Milde-Langosch, K., Loning, T., and Bamberger, C.M. (2001). The glucocorticoid receptor is specifically expressed in the stromal compartment of the human endometrium. J Clin Endocrinol Metab 86, 5071–5074.

Bever, A.T., Hisaw, F.L., and Velardo, J.T. (1956). Inhibitory action of desoxycorticosterone acetate, cortisone acetate, and testosterone on uterine growth induced by estradiol-17beta. Endocrinology 59, 165–169.

Bitman, J., and Cecil, H.C. (1967). Differential inhibition by cortisol of estrogen-stimulated uterine responses. Endocrinology 80, 423–429.

Bolt, M.J., Stossi, F., Newberg, J.Y., Orjalo, A., Johansson, H.E., and Mancini, M.A. (2013). Coactivators enable glucocorticoid receptor recruitment to fine-tune estrogen receptor transcriptional responses. Nucleic Acids Res 41, 4036.

Buenrostro, J.D., Giresi, P.G., Zaba, L.C., Chang, H.Y., and Greenleaf, W.J. (2013). Transposition of native chromatin for fast and sensitive epigenomic profiling of open chromatin, DNA-binding proteins and nucleosome position. Nat Methods 10, 1213–1218.

Cancer Genome Atlas Research, N., Kandoth, C., Schultz, N., Cherniack, A.D., Akbani, R., Liu, Y., Shen, H., Robertson, A.G., Pashtan, I., Shen, R., et al. (2013). Integrated genomic characterization of endometrial carcinoma. Nature 497, 67–73.

Carisey, A., and Ballestrem, C. (2011). Vinculin, an adapter protein in control of cell adhesion signalling. Eur J Cell Biol 90, 157–163.

Citi, S., Sabanay, H., Jakes, R., Geiger, B., and Kendrick-Jones, J. (1988). Cingulin, a new peripheral component of tight junctions. Nature 333, 272–276.

D’Amato, N.C., Gordon, M.A., Babbs, B., Spoelstra, N.S., Carson Butterfield, K.T., Torkko, K.C., Phan, V.T., Barton, V.N., Rogers, T.J., Sartorius, C.A., et al. (2016). Cooperative Dynamics of AR and ER Activity in Breast Cancer. Mol Cancer Res 14, 1054–1067.

Deng, L., Feng, J., and Broaddus, R.R. (2010). The novel estrogen-induced gene EIG121 regulates autophagy and promotes cell survival under stress. Cell Death Dis 1, e32.

Gunin, A.G., Mashin, I.N., and Zakharov, D.A. (2001). Proliferation, mitosis orientation and morphogenetic changes in the uterus of mice following chronic treatment with both estrogen and glucocorticoid hormones. J Endocrinol 169, 23–31.

Haynes, L.E., Lendon, C.L., Barber, D.J., and Mitchell, I.J. (2003). 17 Beta-oestradiol attenuates dexamethasone-induced lethal and sublethal neuronal damage in the striatum and hippocampus. Neuroscience 120, 799–806.

Hertz, G.Z., and Stormo, G.D. (1999). Identifying DNA and protein patterns with statistically significant alignments of multiple sequences. Bioinformatics 15, 563–577.

Isikbay, M., Otto, K., Kregel, S., Kach, J., Cai, Y., Vander Griend, D.J., Conzen, S.D., and Szmulewitz, R.Z. (2014). Glucocorticoid receptor activity contributes to resistance to androgen-targeted therapy in prostate cancer. Horm Cancer 5, 72–89.

Jiao, X., Sherman, B.T., Huang, D.W., Stephens, R., Baseler, M.W., Lane, H.C., and Lempicki, R.A. (2012). DAVID-WS: a stateful web service to facilitate gene/protein list analysis. Bioinformatics 28, 1805–1806.

Jozwik, K.M., and Carroll, J.S. (2012). Pioneer factors in hormone-dependent cancers. Nat Rev Cancer 12, 381–385.

Karmakar, S., Jin, Y., and Nagaich, A.K. (2013). Interaction of glucocorticoid receptor (GR) with estrogen receptor (ER) α and activator protein 1 (AP1) in dexamethasone-mediated interference of ERα activity. J Biol Chem 288, 24020–24034.

Kim, D., Langmead, B., and Salzberg, S.L. (2015). HISAT: a fast spliced aligner with low memory requirements. Nat Methods 12, 357–360.

Knudsen, K.A. (1995). Interaction of alpha-actinin with the cadherin/catenin cell-cell adhesion complex via alpha-catenin. J Cell Biol 130, 67–77.

Lam, K.S., Lee, M.F., Tam, S.P., and Srivastava, G. (1996). Gene expression of the receptor for growth-hormone-releasing hormone is physiologically regulated by glucocorticoids and estrogen. Neuroendocrinology 63, 475–480.

Langmead, B., Trapnell, C., Pop, M., and Salzberg, S.L. (2009). Ultrafast and memory-efficient alignment of short DNA sequences to the human genome. Genome biology 10, R25.

Li, H., Handsaker, B., Wysoker, A., Fennell, T., Ruan, J., Homer, N., Marth, G., Abecasis, G., Durbin, R., and Genome Project Data Processing, S. (2009). The Sequence Alignment/Map format and SAMtools. Bioinformatics 25, 2078–2079.

Li, J., Alyamani, M., Zhang, A., Chang, K.-H., Berk, M., Li, Z., Zhu, Z., Petro, M., Magi-Galluzzi, C., Taplin, M.-E., et al. (2017). Aberrant corticosteroid metabolism in tumor cells enables GR takeover in enzalutamide resistant prostate cancer. Elife 6.

Liao, Y., Smyth, G.K., and Shi, W. (2014). featureCounts: an efficient general purpose program for assigning sequence reads to genomic features. Bioinformatics 30, 923–930.

Love, M.I., Huber, W., and Anders, S. (2014). Moderated estimation of fold change and dispersion for RNA-seq data with DESeq2. Genome biology 15, 550.

Markaverich, B.M., Upchurch, S., and Clark, J.H. (1981). Progesterone and dexamethasone antagonism of uterine growth: a role for a second nuclear binding site for estradiol in estrogen action. J Steroid Biochem 14, 125–132.

Meyer, M.E., Gronemeyer, H., Turcotte, B., Bocquel, M.T., Tasset, D., and Chambon, P. (1989). Steroid hormone receptors compete for factors that mediate their enhancer function. Cell 57, 433–442.

Miranda, T.B., Voss, T.C., Sung, M.-H., Baek, S., John, S., Hawkins, M., Grøntved, L., Schiltz, R.L., and Hager, G.L. (2013). Reprogramming the chromatin landscape: interplay of the estrogen and glucocorticoid receptors at the genomic level. Cancer Res 73, 5130–5139.

Mohammed, H., Russell, I.A., Stark, R., Rueda, O.M., Hickey, T.E., Tarulli, G.A., Serandour, A.A., Serandour, A.A.A., Birrell, S.N., Bruna, A., et al. (2015). Progesterone receptor modulates ERα action in breast cancer. Nature 523, 313–317.

Pan, D., Kocherginsky, M., and Conzen, S.D. (2011a). Activation of the glucocorticoid receptor is associated with poor prognosis in estrogen receptor-negative breast cancer. Cancer Res 71, 6360–6370.

Pan, D., Kocherginsky, M., and Conzen, S.D. (2011b). Activation of the glucocorticoid receptor is associated with poor prognosis in estrogen receptor-negative breast cancer. Cancer Res 71, 6360–6370.

Picelli, S., Björklund, A.K., Reinius, B., Sagasser, S., Winberg, G., and Sandberg, R. (2014). Tn5 transposase and tagmentation procedures for massively scaled sequencing projects. Genome Res 24, 2033–2040.

Quinlan, A.R., and Hall, I.M. (2010). BEDTools: a flexible suite of utilities for comparing genomic features. Bioinformatics 26, 841–842.

Rabin, D.S., Johnson, E.O., Brandon, D.D., Liapi, C., and Chrousos, G.P. (1990). Glucocorticoids inhibit estradiol-mediated uterine growth: possible role of the uterine estradiol receptor. Biol Reprod 42, 74–80.

Reddy, T.E., Pauli, F., Sprouse, R.O., Neff, N.F., Newberry, K.M., Garabedian, M.J., and Myers, R.M. (2009). Genomic determination of the glucocorticoid response reveals unexpected mechanisms of gene regulation. Genome Res 19, 2163–2171.

Rhen, T., Grissom, S., Afshari, C., and Cidlowski, J.A. (2003). Dexamethasone blocks the rapid biological effects of 17beta-estradiol in the rat uterus without antagonizing its global genomic actions. FASEB J 17, 1849–1870.

Robinson, J.L.L., Macarthur, S., Ross-Innes, C.S., Tilley, W.D., Neal, D.E., Mills, I.G., and Carroll, J.S. (2011). Androgen receptor driven transcription in molecular apocrine breast cancer is mediated by FoxA1. EMBO J #1, 3019–3027.

Saso, S., Chatterjee, J., Georgiou, E., Ditri, A.M., Smith, J.R., and Ghaem-Maghami, S. (2011). Endometrial cancer. BMJ 343, d3954.

Singhal, H., Greene, M.E., Tarulli, G., Zarnke, A.L., Bourgo, R.J., Laine, M., Chang, Y.-F., Ma, S., Dembo, A.G., Raj, G.V., et al. (2016). Genomic agonism and phenotypic antagonism between estrogen and progesterone receptors in breast cancer. Sci Adv 2, e1501924.

Tangen, I.L., Werner, H.M., Berg, A., Halle, M.K., Kusonmano, K., Trovik, J., Hoivik, E.A., Mills, G.B., Krakstad, C., and Salvesen, H.B. (2014). Loss of progesterone receptor links to high proliferation and increases from primary to metastatic endometrial cancer lesions. Eur J Cancer 50, 3003–3010.

Terakawa, N., Shimizu, I., Aono, T., Tanizawa, O., and Matsumoto, K. (1985). Dexamethasone suppresses estrogen action at the pituitary level without modulating estrogen receptor dynamics. J Steroid Biochem 23, 385–388.

Voss, T.C., Schiltz, R.L., Sung, M.-H., Yen, P.M., Stamatoyannopoulos, J.A., Biddie, S.C., Johnson, T.A., Miranda, T.B., John, S., and Hager, G.L. (2011). Dynamic exchange at regulatory elements during chromatin remodeling underlies assisted loading mechanism. Cell 146, 544–554.

Westin, S.N., Broaddus, R.R., Deng, L., McCampbell, A., Lu, K.H., Lacour, R.A., Milam, M.R., Urbauer, D.L., Mueller, P., Pickar, J.H., et al. (2009). Molecular clustering of endometrial carcinoma based on estrogen-induced gene expression. Cancer Biol Ther 8, 2126–2135.

Wik, E., Raeder, M.B., Krakstad, C., Trovik, J., Birkeland, E., Hoivik, E.A., Mjos, S., Werner, H.M., Mannelqvist, M., Stefansson, I.M., et al. (2013). Lack of estrogen receptor-alpha is associated with epithelial-mesenchymal transition and PI3K alterations in endometrial carcinoma. Clin Cancer Res 19, 1094–1105.

Yang, C.-H., Almomen, A., Wee, Y.S., Jarboe, E.A., Peterson, C.M., and Janát-Amsbury, M.M. (2015). An estrogen-induced endometrial hyperplasia mouse model recapitulating human disease progression and genetic aberrations. Cancer Med 4, 1039–1050.

Yang, F., Ma, Q., Liu, Z., Li, W., Tan, Y., Jin, C., Ma, W., Hu, Y., Shen, J., Ohgi, K.A., et al. (2017). Glucocorticoid Receptor:MegaTrans Switching Mediates the Repression of an ERalpha-Regulated Transcriptional Program. Mol Cell 66, 321–331 e326.

Zhang, Y., Liu, T., Meyer, C.A., Eeckhoute, J., Johnson, D.S., Bernstein, B.E., Nusbaum, C., Myers, R.M., Brown, M., Li, W., et al. (2008a). Model-based analysis of ChIP-Seq (MACS). Genome Biol 9, R137.

Zhang, Y., Liu, T., Meyer, C.A., Eeckhoute, J., Johnson, D.S., Bernstein, B.E., Nusbaum, C., Myers, R.M., Brown, M., Li, W., et al. (2008b). Model-based analysis of ChIP-Seq (MACS). Genome biology 9, R137.

Zhou, F., Bouillard, B., Pharaboz-Joly, M.O., and André, J. (1989). Non-classical antiestrogenic actions of dexamethasone in variant MCF-7 human breast cancer cells in culture. Mol Cell Endocrinol 66, 189–197.

